# Photorealistic reconstruction of visual texture from EEG signals

**DOI:** 10.1101/2021.08.01.449562

**Authors:** Suguru Wakita, Taiki Orima, Isamu Motoyoshi

## Abstract

Recent advances in brain decoding have made it possible to classify image categories based on neural activity. Increasing numbers of studies have further attempted to reconstruct the image itself. However, because images of objects and scenes inherently involve spatial layout information, the reconstruction usually requires retinotopically organized neural data with high spatial resolution, such as fMRI signals. In contrast, spatial layout does not matter in the perception of ‘texture’, which is known to be represented as spatially global image statistics in the visual cortex. This property of ‘texture’ enables us to reconstruct the perceived image from EEG signals, which have a low spatial resolution. Here, we propose an MVAE-based approach for reconstructing texture images from visual evoked potentials measured from observers viewing natural textures such as the textures of various surfaces and object ensembles. This approach allowed us to reconstruct images that perceptually resemble the original textures with a photographic appearance. A subsequent analysis of the dynamic development of the internal texture representation in the VGG network showed that the reproductivity of texture rapidly improves at 200 ms latency in the lower layers but improves more gradually in the higher layers. The present approach can be used as a method for decoding the highly detailed ‘impression’ of sensory stimuli from brain activity.

## Introduction

In the field of neuroscience, an increasing number of studies have been conducted to estimate perceptual content and psychological states by extracting certain statistical patterns from brain activity data (Kamitani & Tong, 2005; Schwartz et al., 2006; Miyawaki et al., 2008; Carlson et al., 2011; Green & Kalaska, 2011; Nishimoto et al., 2011). A number of ‘brain decoding’ techniques that identify the object category of an image from the fMRI-BOLD signal have been reported (Shenoy & Tan, 2008; Das et al., 2010; Wang et al., 2012; Carlson et al., 2013; Stewart et al., 2014; Kaneshiro et al., 2015). In recent years, ambitious attempts have been made to reconstruct the image itself from brain activity (Palazzo et al., 2017; Shen et al., 2019a; Shen et al., 2019b). For instance, Shen et al. (Shen et al., 2019a) proposed a method of decoding visual features for each hierarchical stage of visual information processing from an fMRI signal using a deep neural network (DNN) (Krizhevsky et al., 2012; Simonyan & Zisserman 2015, He et al., 2016) and successfully reconstructed not only the presented image but also the image that an observer imagined in her/his mind.

While excellent decoding is supported by the big data of fMRI, the scope of application is limited by the high costs and potential invasiveness of fMRI. To overcome this limitation, several studies adopted EEG, which provides an easy, cheap, and non-invasive way to collect brain activity data. Palazzo et al. (Palazzo et al., 2017) introduced a method for reconstructing the image of an object from EEG signals by converting the EEG signals into features and conditioning generative adversarial networks (GANs) (Goodfellow et al., 2014) by it. This approach allowed them to reconstruct an image that can be correctly classified into the original object category (EEG classification accuracy: 84%, Inception Score(IS): 5.07, Inception Classification accuracy(IC): 0.43). However, as pointed out by the authors themselves, their result is a product of a generative model conditioned by categorical information extracted from an EEG signal and not the direct reconstruction of the image itself actually given to the observer. It is evident that this method fails to reproduce aspects of the perceptual realism of an image, such as the detailed shape, sharp contours, and textures. This limitation seems unavoidable considering the small data size of EEG signals, especially in terms of spatial resolution.

Against the above background, it is of interest to explore the use of ‘texture’ images in decoding from EEG signals. The perception of a texture is based on spatially global image statistics (Julesz, 1965; Heeger & Bergen, 1995; Portilla & Simoncelli, 2000; Landy & Graham, 2004; Freeman & Simoncelli, 2011), and it is even possible to synthesize perceptually similar texture images using only those statistics (Portilla & Simoncelli, 2000). Such statistical information is represented in the low- and mid-level visual cortex, such as V1, V2, and V4 (Freeman et al., 2013; Okazawa et al.,2015; Okazawa et al., 2017; Ziemba et al., 2019), and used in the rapid perception of scenes, objects, and surface materials (Thorpe et al., 1996; Oliva & Torralba, 2001; Motoyoshi et al., 2007; Rosenholtz et al., 2012; Whitney et al., 2014). In convolutional neural network (CNN), which computationally mimics neural processing in the ventral stream of the visual brain, the spatially global information obtained by the Gram matrix transformation of features extracted from each hierarchical layer stage corresponds to texture representation (Gatys et al., 2015; Gatys et al., 2016).

According to these findings, it is expected that texture can be reconstructed from EEG signals by estimating the information that correlates with the spatially global statistics for texture representation. In fact, the recent study (Orima & Motoyoshi, 2021) were able to estimate lower-order image statistics from VEPs using a linear regression model and synthesize the texture images with identical image statistics. Using the Image-VEP dataset collected in that study, the present paper proposes a CNN-based method that allows a high quality of reconstruction of the original texture image from a VEP for a variety of natural textures.

## Method

Texture perception is essentially based on the visual appearance, or impression, of an image according to the continuous perceptual similarity, rather than categorical conceptual knowledge as required for object recognition. From this view, we specifically adopted an MVAE-based approach (Suzuki et al., 2017; Wu & Goodman, 2018; Kurle et al., 2019; Shi et al, 2019; Tsai et al, 2019) that acquires a continuous latent representation shared by a texture image and EEG signal. Using the trained MVAE model, we attempted to reconstruct the texture image from the latent variables obtained when only one-modality information, EEG data, was input.

In our approach, the MVAE model is trained with the texture images and VEP as two-modality information. After training, the latent space shared by the two modalities is acquired in the model. Finally, the test texture image is reconstructed from the latent variable obtained from the corresponding EEG signals input to the trained model.

### EEG measurement

In training the model, we used the dataset obtained by Orima & Motoyoshi (2021). The dataset comprises EEG signals for 166 natural texture images, with each signal measured for a period of 500 ms, 24 times, for each of 15 human observers.

**Figure 1.**
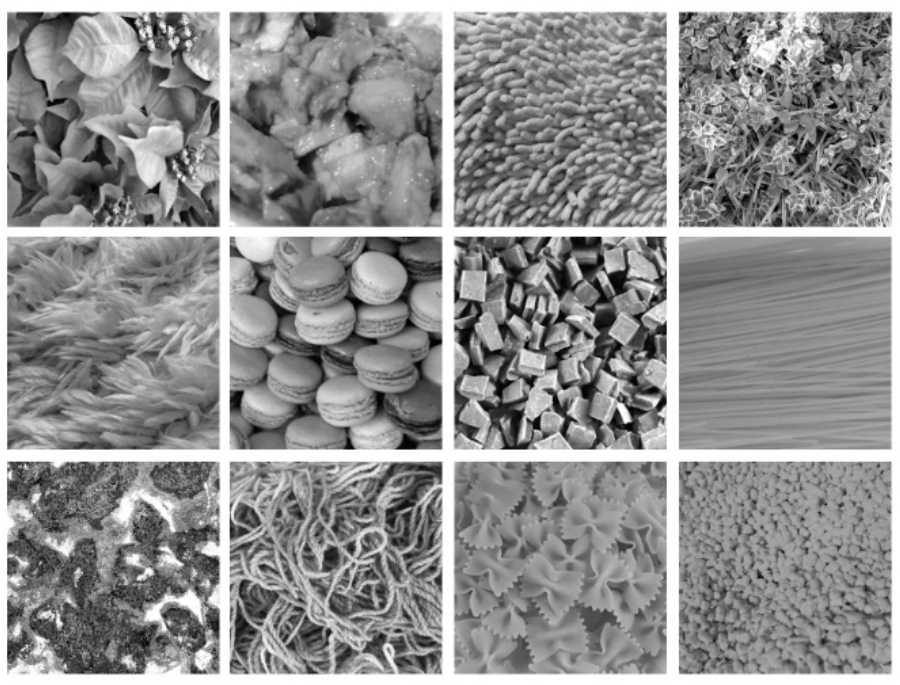
Examples of texture images used in the EEG measurement.

Visual stimuli were images of 166 natural textures subtending 5.7 deg × 5.7 deg (256 × 256 pixels). The images were collected from the Internet and our own image database. Each image was achromatic and had a mean luminance of 33 cd/m^2^. In each of 24 measurement blocks, 166 images were presented in random order for 500 ms followed by a 750-ms blank that is equal to a uniform gray background and 15 observers viewed each image with their eyes steadily fixed at the center of the image. During each block, the VEP was measured using 19 electrodes (Fp1, Fp2, F3, F4, C3, C4, P3, P4, O1, O2, F7, F8, T7, T8, P7, P8, Fz, Cz, and Pz according to the international 10/20 method; BrainVision Recorder, BrainAmp Amplifier, EasyCap; Brain Products GmbH). All stimuli were presented on a gamma-corrected LCD (BENQ XL2420T). The refresh rate of the LCD was 60 Hz, and the spatial resolution was 1.34 min/pixel at an observation distance of 100 cm. All measurements were conducted in accordance with the Ethics Committee for Experiments on Humans at the Graduate School of Arts and Sciences, The University of Tokyo. The participants completed a written consent form.

### MVAE for image reconstruction from EEG signals

Considering the continuous and variegated nature of natural textures as visual information, we consider a variational auto encoder (VAE) -based (Kingma & Welling, 2013) approach in which the texture images and the corresponding EEG signals are represented in a continuous latent space.

The VAE is a deep generative model that conducts its generation process by deep learning assuming the existence of a latent variable *z* when data *v* are observed (Kingma & Welling, 2013; Kingma et al., 2014; Krishnan et al., 2015; Dai et al., 2015). Here, by assuming that latent variables are represented on a probabilistic distribution space, we can perform continuous representation learning on the observed input data (Equation 1).

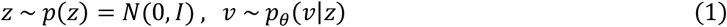

In the VAE, the observed input data *v* are transformed by the encoder into a contractive intermediate representation called latent variable *z*, and the decoder reconstructs the original input data *v*′ with this latent variable as input. The entire model is trained so as to minimize the difference between the input data *v* and the reconstructed data *v*′, and the model parameters of the encoder and decoder are updated. The encoder and decoder comprise a neural network. (In the following, *θ* and *Φ* refer to the model parameters of the encoder and decoder, respectively, and the multivariate Gaussian distribution is denoted p(z).)

More practically, the target of training is to maximize the marginal likelihood *p_θ_*(*x*), but because this cannot be treated directly, we optimize the model parameters of the encoder *q_ϕ_*(*z*|*x*) and decoder *p_θ_*(*x*|*z*) to maximize the evidence lower bound (ELBO) given in Equation 2.

In Equation 2, the first term on the right-hand side is called the regularization term. This term regularizes the latent variable z, which is obtained basing on the mean vector *μ* and variance vector *σ* output by the encoder, to distribute according to prior *p*(*z*).

The second term on the right-hand side is the reconstruction error term, which minimizes the difference between the original input data *v* and *v*′, the input data reconstructed from the decoder using the latent variable z.

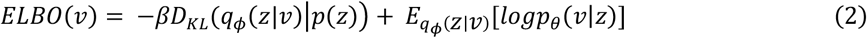

As an extension of the VAE, the multimodal VAE, which treats multimodal information as input, has been proposed (Suzuki et al., 2017; Wu & Goodman, 2018; Kurle et al., 2019; Shi et al, 2019; Tsai et al, 2019). This extension is inspired by the fact that our cognition in the real world uses multimodal information, not unimodal information (Ngiam et al.,2011; Srivastava & Salakhutdinov, 2012; Kiros et al., 2014; Pandey & Dukkipati, 2016). In fact, it is generally known that learning with multimodal information induces the acquisition of better informative representations compared with the case of unimodal information (Ngiam et al., 2011; Srivastava & Salakhutdinov, 2012).

In this study, we apply the extended method for the MVAE (Wu & Goodman, 2018), which allows inference of latent variables even under the partial observation of multimodal information aiming at reconstructing texture images only from EEG signals.

Here, the texture images and EEG signals are treated as different information modalities, and the latent representation shared by these two modalities is acquired by the learning MVAE. As a result of this training, the stimulus can be reconstructed by decoding the texture image using latent variables acquired by the input of a single modality, the EEG signal. Figure 2 is an overview of the structure of the MAVE model. The MVAE model comprises an encoder and decoder for the EEG signal modality and an encoder and decoder for the texture image modality. By inputting one or both of the modalities of information into the encoder corresponding to the respective modal information, a latent variable can be inferred. The latent variables obtained here are integrated into a single latent variable using the product of experts (PoE) (Hinton, 2002). Finally, the reconstructed results of EEG signals and texture images are obtained by inputting this latent variable to each of the decoders corresponding to each modal information.

**Figure 2.**
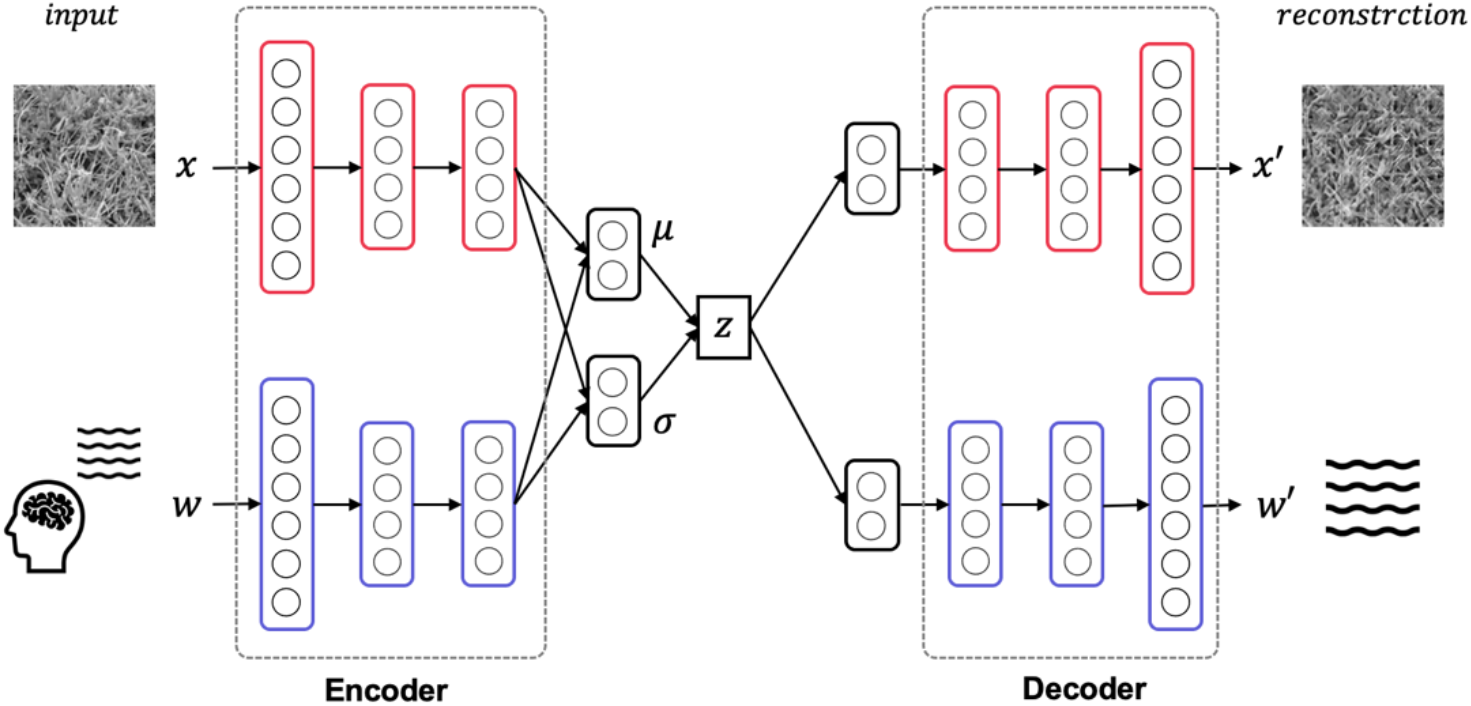
Model overview. The MVAE model comprises an encoder and decoder for the EEG signal modality and an encoder and decoder for the texture stimulus modality. After training this model, the latent space shared by the two modalities is acquired. Thus, the texture stimulus that was presented in the EEG measurement can be reconstructed from the latent variable obtained by inputting only the EEG signals to this trained model.

In the training of the MVAE model, there are three possible patterns for the combination of the observable modal information. (Here, we denote the texture image modal information as *x*, the EEG signal modal information as *w*, and the observed whole or partial modal information as *V* = {*v*: *v*_1_, *v*_2_,…*v_n_*}}.)

- *V*_1_ = {*v*:*x,w*}: Both modalities, the EEG signal and texture image, can be observed.
- *V*_2_ = {*v*:*x*}: One modality, the texture image, can be observed.
- *V*_3_ = {*v*:*w*}: One modality, the EEG signal, can be observed.

In reconstructing texture images from EEG signals, which is the target of this study, it is necessary to obtain a representation in the latent space shared by the two modalities of texture images and EEG signals, and to be able to extract latent variables under the partial modal observation (*V*_2_, *V*_3_) that are as good as or similar to those extracted under the full modal observation (*V*_1_). Considering this point, we maximize the ELBO expressed in Equation 3, proposed by (Wu & Goodman, 2018), in our training.

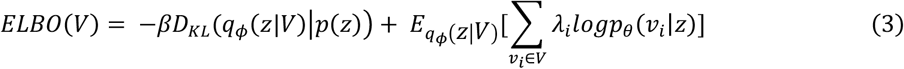

The loss function of the entire model is thus expressed by Equation 4, and the training proceeds accordingly.

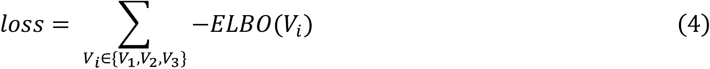

One issue that should be considered here is that the image reconstructed using the VAE-based approach is generally blurred. When we tested the reconstruction with the simple VAE using texture images, we found that the reconstruction of fine texture components did not work well, resulting in grayish or blurred images. This is a crucial issue in the present study because we are aiming to realize texture reconstruction with visual similarity to the texture stimuli presented to the observers during EEG measurement. As a solution to such problems, a method combining a VAE and GAN (Rosca et al., 2017) has been proposed to generate natural images and general object images more realistically. However, in the present paper, it is necessary to devise a loss function that improves the reproduction for such texture components when we consider that we use natural texture images in the present study and particularly when failing to reconstruct fine and relatively high spatial frequency components. We thus considered applying precedent knowledge gained in the field of neural style transfer (Gatys et al., 2016; Johnson et al., 2016; Ulyanov et al., 2016; Huang & Belongie, 2017), where texture synthesis is conducted using trained deep neural network (Gatys et al., 2015). According to this knowledge, in the trained VGG-19 model (Simonyan & Zisserman, 2015), style information at different levels of abstraction is processed at each stage of the hierarchical processing, and we can extract fine style information at the lower layers and global style information at the higher layers. Additionally, this style information is sufficient for accurate style transferring and texture synthesis (Gatys et al., 2015; Gatys et al., 2016; Johnson et al., 2016; Ulyanov et al., 2016; Huang & Belongie, 2017). We therefore use this style information in our approach for more precise texture reconstruction. Specifically, we replace the reconstruction error term in the ELBO with a combination of original reconstruction error term and style error term, which is commonly used in the framework of neural style transfer. The style error is expressed in Equation 5. Here, we denote the input image as *x*, reconstructed image as *x*′, and set of layers in the trained VGG-19 from which the style information can be extracted as L = {1,2, … k}. Style information obtained by Gram matrix transformation of the output from each layer is denoted 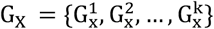 and 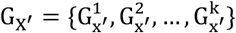 for the input image and reconstructed image respectively. α is the weighting of style information in each layer, N is the number of filter maps in each layer of VGG-19, and M is the number of elements in each filter map in each layer.

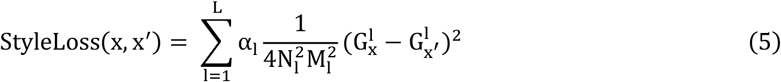

Applying this style error for the loss function confirmed that the texture pattern can be reconstructed clearly regardless of the spatial frequency of the texture in the input image.

## Results

### MVAE model and training

Following the previous EEG decoding study (Palazzo et al., 2017), we used all 166 texture images and corresponding EEG signals in training the model. Instead, for each texture image, the corresponding EEG signals were divided into training, validation, and test datasets according to the proportions of 80%, 10%, and 10% for training the MVAE model. The EEG signals, which were measured for 500 ms after the stimulus onset when the texture stimulus was presented to the observer, were treated as raw vector data with 500 dimensions. When we input EEG signals to the MVAE model, 25–30 samples of EEG signals corresponding to one particular texture stimuli were selected in random combinations and their average waveforms were normalized in the range of min0 to max1. Among the electrode channels used in the EEG measurement, the signals measured at Fp1, Fp2, F3, F4, C3, C4, P3, P4, O1, O2, F7, F8, T7, T8, P7, and P8 in the international 10/20 method were used as input. Additionally, texture images, as the other information modality, was resized to 128 × 128 on input. At this time, the reconstructed texture image was also output as a 128 × 128 image. In the training, the Adam gradient descent method was used with a learning rate of 1e–4. The batch size was 16. The vector size for the latent variable of the MVAE model was 256. The MVAE model comprises an encoder and decoder that treat the texture images as modal information and an encoder and decoder that treat the EEG signals as modal information. We used 2D-convolution for the encoder and decoder that treat the texture images as modal information, and 1D-convolution for the encoder and decoder that treat the EEG data as modal information. The architectural details of the MVAE model are given in Table 1. In actual implementation, except for the final output layer, each convolution layer is followed by batch normalization and ReLU rectifier processing in order.

**Table 1.**
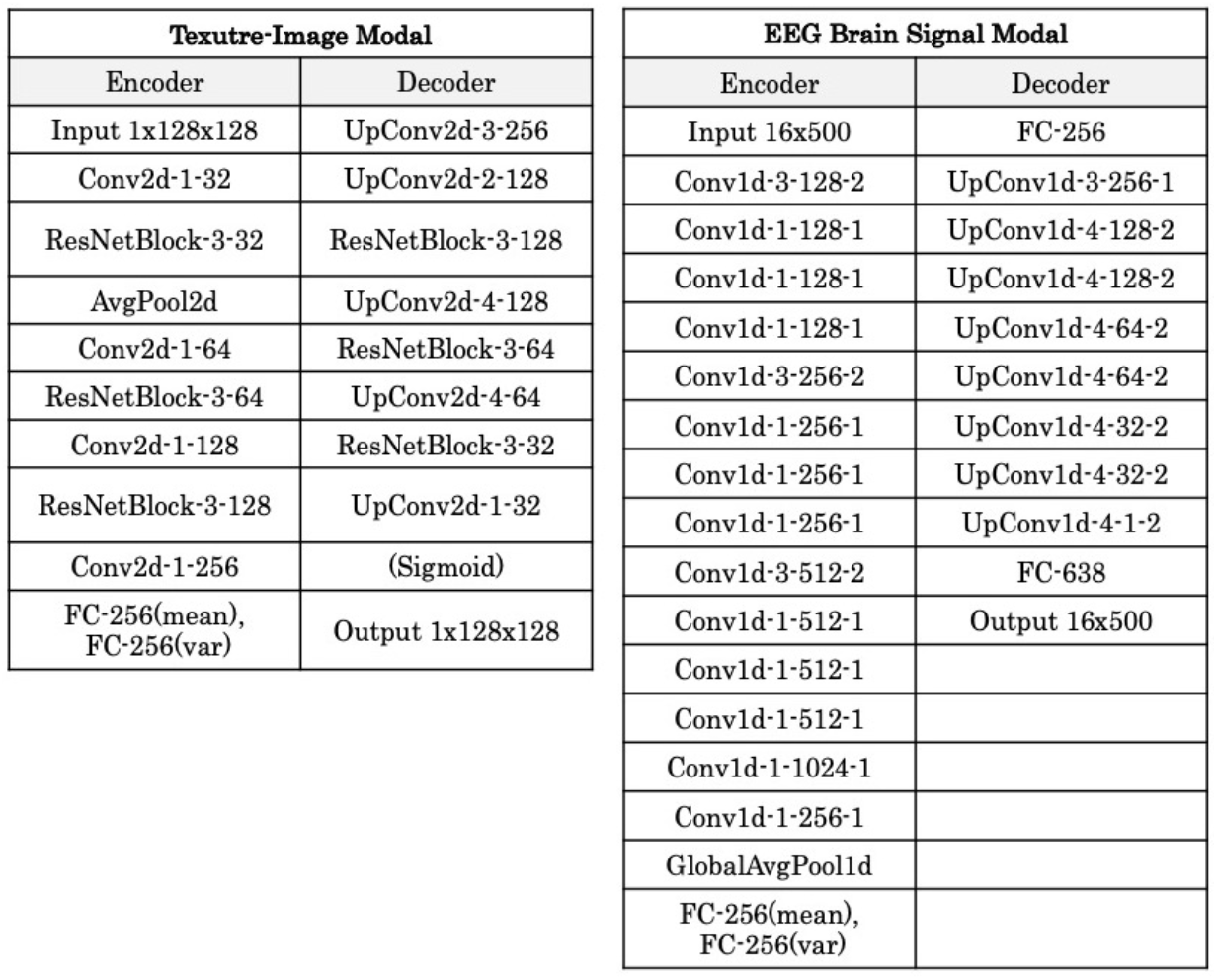
Details of the model architecture

### Reconstruction of the texture image

After training the MVAE model, we reconstructed the texture image using the test EEG signals as input. More specifically, the latent variables were extracted from the encoder that treats the EEG signals as modal information, and the texture images were reconstructed by inputting these latent variables to the other decoder that treats the texture image as modal information.

Figure 3 shows examples of reconstructed images. In each row, the upper images show the original textures and the lower images show the textures reconstructed from EEG. It is seen that most of the reconstructed textures are remarkably photorealistic, and some are similar to the original textures. The quality of reconstruction is much higher than that of texture synthesis based on linear regression reported in our previous study (Orima & Motoyoshi, 2021).

**Figure 3.**
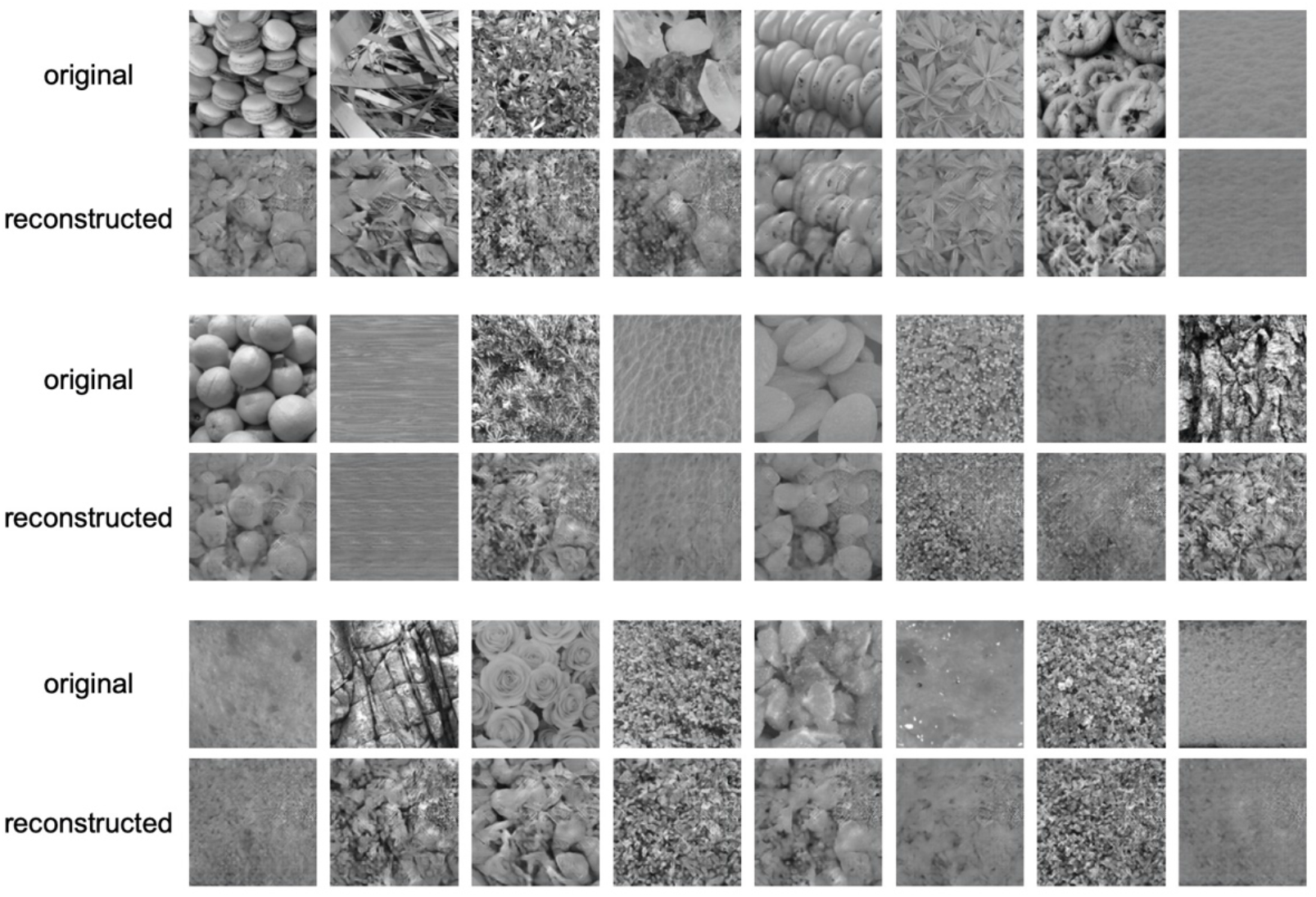
Examples of reconstructed texture images. In each row, the upper images are the original texture images shown to the observer and the lower images are the images reconstructed from EEG.

### Psychophysical experiment

In validating the reconstruction results, we carried out a behavioral experiment to examine the relative perceptual similarity of the reconstructed texture to the original. In our display, the original natural texture (2.6 × 2.6 deg, 128 × 128 pixels) was presented at the center, and reconstructed textures were presented on the left and right, 3.5 deg from the center. One reconstructed texture was the target image reconstructed from EEG signals for the central original texture, and the other was the non-target image reconstructed from EEG signals for another texture that was chosen randomly from a set of 165 textures. Six observers with normal or corrected-to-normal vision viewed the stimuli with a free gaze and indicated the texture image (left/right) that was perceptually more similar to the central original texture. To prevent pixelwise matching and the use of memory, each reconstructed texture was randomly chosen from three possible samples. Observers were strongly instructed to evaluate the similarity in terms of the visual appearance and not in terms of the categorical meaning. For each observer, at least four data were collected and the probability of a response that “the target appeared more similar” was calculated for each image. All experiments were conducted using gamma-corrected LCDs with a 60-Hz refresh rate (SONY PVM 2541A, SONY PVM-A250, BenQ XL2730Z, SONY PVM-A250, BENQ XL2720B, and BENQ XL 2430T), each of which was installed in a dark room of the individual observer’s home. The viewing distance was adjusted so that the spatial resolution was 1.0 min/pixel. Other parameters were the same as those in the EEG measurements.

Figure 4 shows the probability of a response that “the target appeared more similar” averaged across six observers for 166 textures. The horizontal axis is the index of the texture image, sorted from the left in descending order of the proportion correct. A horizontal red line denotes the chance level (50%). For 78.1% of textures, the target image was chosen with a probability significantly higher than the chance level. Together with the observations presented in Figure 3, these results suggest that the reconstruction was fairly successful.

**Figure 4.**
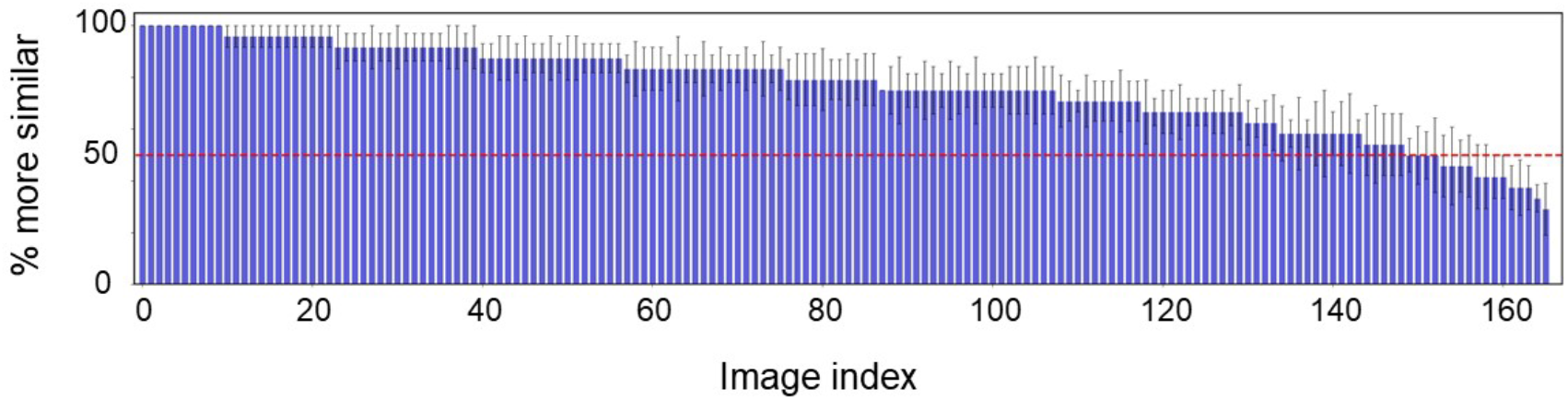
Probability of a response that the image reconstructed from the original texture (target) was more similar than the image reconstructed from another random texture (non-target) to the original image. The horizontal axis shows the index of the texture image, sorted from the left in descending order of the probability. The red line denotes the chance level (50%). The error bars indicate ±1 s.e.m. across observers.

### Temporal development of the texture representation

In investigating how the neural representations corresponding to the texture representation that enables such reconstructions evolve over time, we reconstructed texture images from EEG signals with various temporal periods from the stimulus onset and compared their neural representations with those obtained for the entire interval (i.e., 500 ms).

For this analysis, we prepared 10 samples of EEG signals for each of the 166 original textures of the test dataset used in training the MVAE model. We then input the EEG signals for 100 different temporal periods increasing in length in 5-ms increments from 0 ms (i.e., 0–5 ms, 0–10 ms, … 0–500 ms) into the trained MVAE model, and reconstructed the texture image using only the texture representation included in the EEG signal in each period. We denote the image reconstructed from the EEG signal in the entire interval of 0–500 ms as *x^recon^* and the texture images reconstructed from the EEG signal in each recon period as 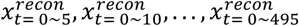.

Figure 5 illustrates the texture images reconstructed with five different temporal periods of the EEG signal. The rightmost image shows the original texture. It is seen that the reconstruction becomes more sophisticated as the temporal period increases.

**Figure 5.**
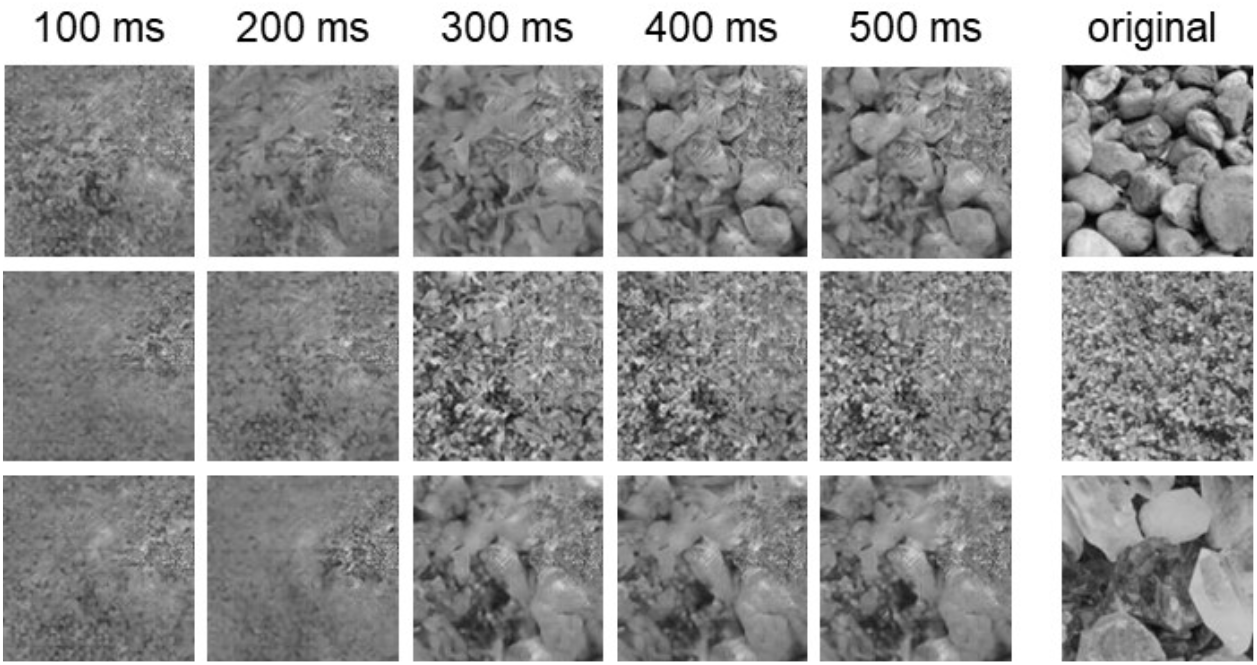
Development of reconstructed texture images over time. In each row, texture images are reconstructed from EEG signals with temporal periods ranging from 100 to 500 ms (total duration). The rightmost image shows the original texture.

We quantitatively evaluated the reproducibility of texture in the texture images reconstructed from EEG signals in each period using the texture representation metric. (Texture representation is referred to as style information in section 2. Method, but we use the term *texture representation* in discussing texture here.) Texture representation can be extracted at each layer stage of the trained VGG, which is commonly used in the field of neural style transfer (Equation 5). Here, we input the image reconstructed from the EEG signal in a certain period 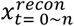 into the trained VGG model and extracted texture representation from the five layers, from the lower layer to upper layer. We conducted the same extraction process for the image reconstructed from the EEG signal including the entire time interval (0–500 ms), *x^recon^*. The difference in texture representation between 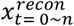 and *x^recon^* was then calculated in terms of the root mean square (RMS) for each of the five layers. The calculation was performed for each EEG signal and each period n, which was from 5 to 500 ms. Finally, values were averaged within each layer.

Figure 6 plots the development of texture representation over time for each layer of the VGG (denoted by color). The horizontal axis represents the temporal period of the EEG signal, and the vertical axis represents the RMS difference in texture representation between the image reconstructed for that period and the image reconstructed for the entire period (0–500 ms); the difference is normalized to the value for the shortest period (0–5 ms). It is assumed that a smaller relative difference in texture representation indicates better reproducibility of the texture representation.

**Figure 6.**
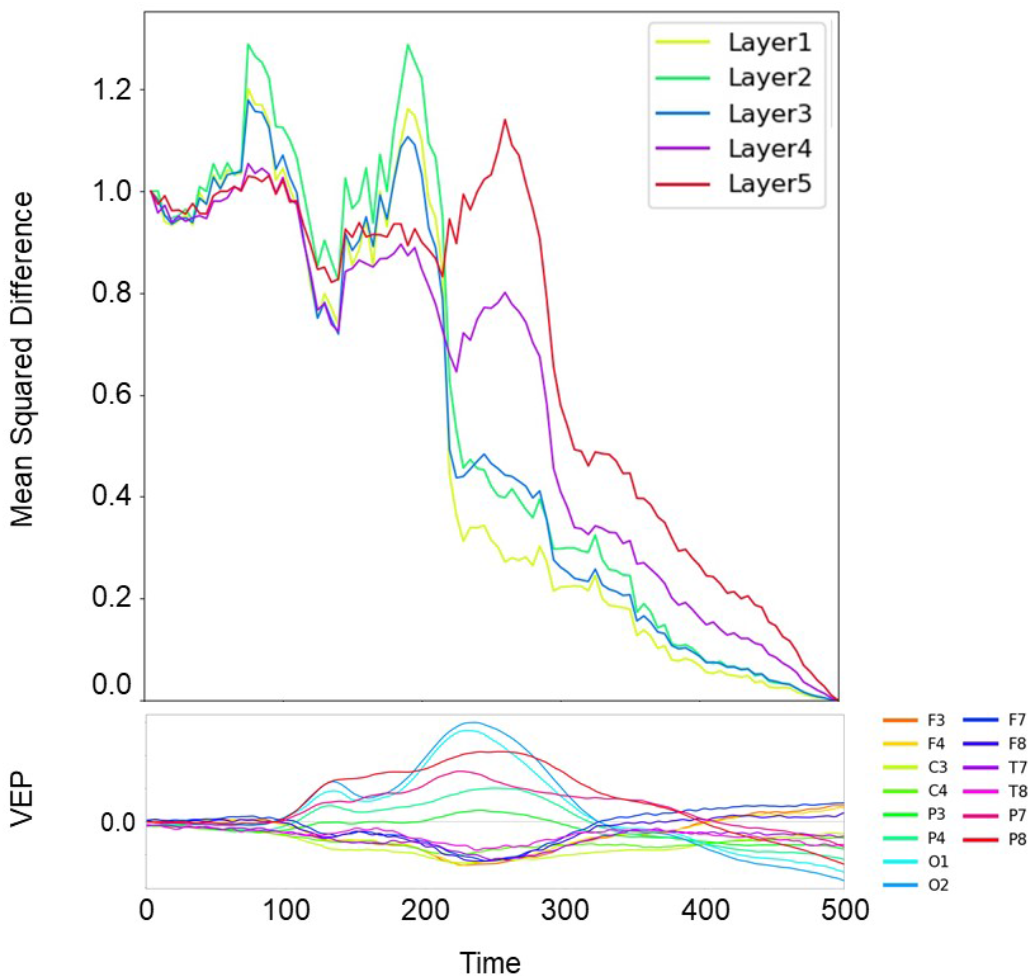
Temporal change in the texture representation of the reconstructed image in each layer of the VGG. The horizontal axis represents the temporal period of the EEG, and the vertical axis represents the normalized RMS difference of texture representation from that for the entire period (0–500 ms). Different colors represent the results for different layers of the VGG. The lower panel shows the grand average VEPs for each electrode.

The results indicate that the reproducibility of texture representation increases with the temporal period of EEG signals. It is seen that, in the lower layers, namely layers 1, 2, and 3, the reproducibility abruptly increases immediately after 200 ms. This is consistent with the time in which VEPs are correlated with the Portilla–Simoncelli texture statistics (Portilla & Simoncelli, 2000; Orima & Motoyoshi, 2021) that are represented in V1 and V2 of the primate brain (Freeman et al., 2013). In the higher layers, namely layers 4 and 5, the reproducibility appears to gradually increase especially after 300 ms, suggesting that high-level texture representation develops slowly.

## Discussion

The present study introduced a method in which an MVAE is used to reconstruct the image of a natural texture from EEG signals alone. Our trained MVAE model successfully reconstructed the original texture with photorealistic quality and greatly outperformed linear regression on the same dataset (Orima & Motoyoshi, 2021). Analysis of the reproducibility of texture representation with temporal accumulation of the EEG signal revealed that the texture representation of the higher layers converges more slowly than that of the lower layers.

As mentioned earlier, it is generally challenging to decode neural representations of a natural scene with EEG because of the low retinotopic resolution of EEG as compared with that of fMRI. The present study avoided this limitation by confining the scope to textures to textures for which the perception is determined by spatially global image statistics, and we successfully reconstructed various natural textures from EEG signals. The previous study having a similar scope (Orima & Motoyoshi, 2021) focused on understanding the neural dynamics for image statistics assumed in human texture perception (e.g., Portilla–Simoncelli statistics) and demonstrated a reconstruction of textures using image statistics linearly regressed from EEG signals. In contrast, the present study pursued a technique to reconstruct an image with higher quality and showed that the use of an MVAE allows the reconstruction of textures with high quality.

The previous approach reconstructed natural object images from EEG signals on the basis of the classification of discrete object categories acquired in a supervised network (Palazzo et al., 2017). However, the resulting images did not look naturalistic even though they were classified into the correct object category. In contrast, the present study aimed to reconstruct a purely perceptual impression without any dependency on top-down knowledge such as that of categories, by acquiring a continuous representation space of visual textures in a fully unsupervised learning manner. The resulting images duplicated the perceptual impression well. Of course, such success might be possible only for the textures that we used, and it is unclear if the present approach is applicable to a wide range of classes of images, such as images of objects and scenes. However, we believe that the fact that we were able to reproduce images from EEG signals in a highly realistic manner brings a new direction in the decoding of sensory information. We are currently applying the same approach to sounds.

We should also note a limitation of the present approach. Figure 7 shows the worst examples of texture reconstruction. The upper images show the original texture, and the other images show the image samples reconstructed from EEG signals. These reconstructed images are similar to one of the other textures tested, and there is a large variability among samples for the same original texture. This result is due to the VAE acquiring continuity on visual similarity between the considered textures in the latent space, and therefore, when the proper texture representation was not extracted from the EEG signals, the representation became an intermediate representation that was determined virtually randomly in the space defined by the limited number of textures that we used. As a result, it is highly possible that the reconstructed texture is similar to one of the other original textures. This problem could be avoided with a latent space that is richer with more diverse texture images. However, such a latent space would require many more images and corresponding EEG data.

**Figure 7.**
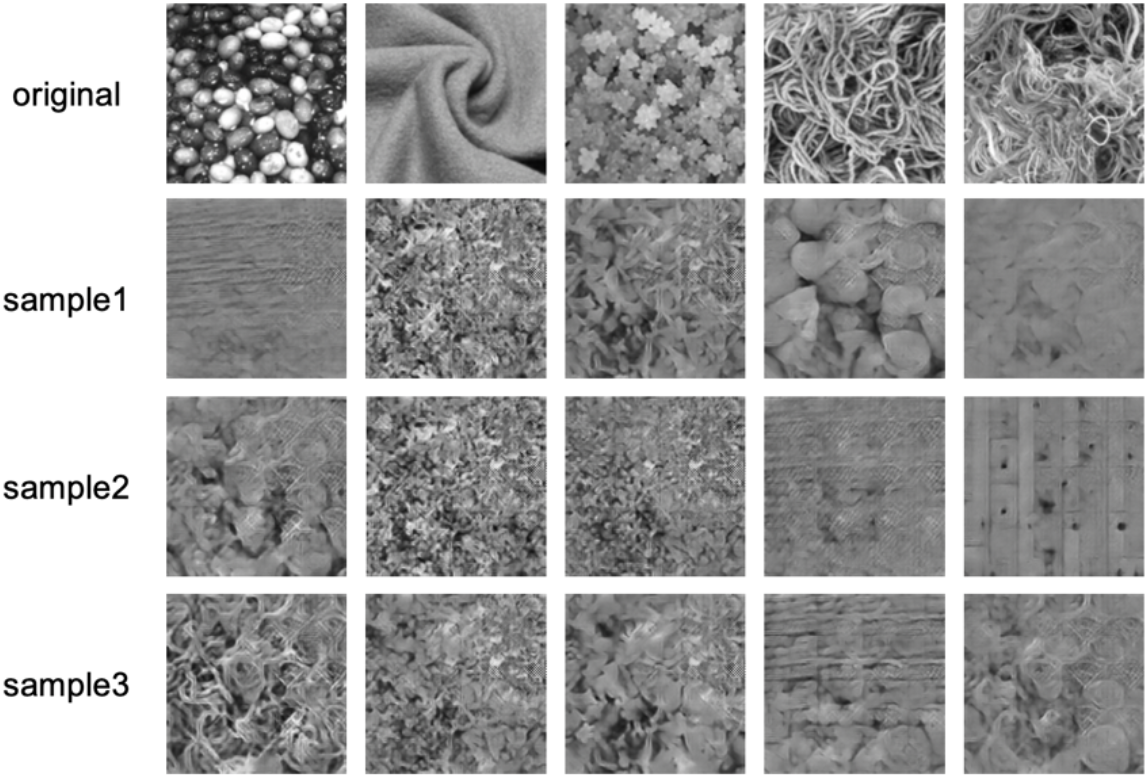
Failure examples of the reconstructed texture images having the lowest ‘more similar’ response rates in the psychophysical experiment. The upper row shows the original textures and the other rows show three sample images reconstructed from EEG signals.

Analysis on the dynamics of texture representation with temporal accumulation in the EEG signal showed that texture representation becomes sophisticated quickly in the lower layers of the VGG and more slowly in the higher layers of the VGG. In the lower layers, the texture representation converges rapidly from a noisy state around 200 ms after the stimulus onset. This timing coincides with the temporal epoch (200–250 ms) in which texture VEPs are strongly correlated with Portilla–Simoncelli image statistics especially at high spatial frequencies and with high-level information beyond image statistics (Orima & Motoyoshi, 2021). In contrast, the texture representation in higher layers converges slowly beyond 300 ms after the onset. This seems to be related with high-level cortical representations of global pattern information that integrates the output of the early visual cortex over a large spatial range (Gallant et al., 1993; Schyns & Oliva, 1994; Okazawa et al., 2015; Portilla & Simoncelli, 2000; Pasupathy et al., 2020). The other possible interpretation is that this sluggish development reflects changes in neural representation thorough the feedback loop in the visual cortex (Lamme et al., 1998; Bondy et al., 2018; Liang et al., 2017).

While the present approach provides an effective tool for reconstructing the visual impression of an image with complex spatial structures from EEG signals, there is still room for improvement. By applying frameworks such as the conditional VAE, the present work is expected to be extended to the analysis of visual impressions specific to a particular individual and the examination of particular visual functions. In the present study, we focused on the pipeline of reconstructing texture stimuli from EEG signals, but owing to the nature of MVAE-based systems, it is also possible to consider the opposite pipeline through which the EEG signal is reconstructed from an image.

## Acknowledgment

This study was supported by Commissioned Research of NICT (1940101) and JSPS KAKENHI JP20K21803.

